# The genomic and transcriptomic landscape of advanced renal cell cancer for individualized treatment strategies

**DOI:** 10.1101/2022.04.22.488773

**Authors:** K. de Joode, W.S. van de Geer, G.J.L.H. van Leenders, P. Hamberg, H.M. Westgeest, A. Beeker, S.F. Oosting, J.M. van Rooijen, L.V. Beerepoot, M. Labots, R.H.J. Mathijssen, M.P. Lolkema, E. Cuppen, S. Sleijfer, H.J.G. van de Werken, A.A.M. van der Veldt

## Abstract

**Background:** Differences in the clinical course and treatment responses in individual patients with advanced renal cell carcinoma (RCC) can largely be explained by the different genomics of this disease. To improve the personalized treatment strategy and survival outcomes for patients with advanced RCC, the genomic make-up in patients with advanced RCC was investigated to identify putative actionable mutations and signatures.

**Methods:** In this prospective multicenter study (NCT01855477), whole-genome sequencing (WGS) data of locally advanced and metastatic tissue biopsies and matched whole-blood samples were collected from 91 patients with histopathologically confirmed RCC. WGS data were analyzed for small somatic variants, copy-number alterations and structural variants. For a subgroup of patients, RNA sequencing (RNA-Seq) data could be analyzed. RNA-Seq data were clustered on immunogenic and angiogenic gene expression patterns according to a previously developed angio-immunogenic gene signature.

**Results:** For papillary and clear cell RCC, putative actionable drug targets were detected by WGS in 100% of the patients. RNA-Seq data of clear cell and papillary RCC were clustered using a previously developed angio-immunogenic gene signature. Analyses of driver mutations and RNA-Seq data revealed clear differences among different RCC subtypes, showing the added value of WGS and RNA-Seq over clinicopathological data.

**Conclusions:** By improving both histological subtyping and the selection of treatment according to actionable targets and immune signatures, WGS and RNA-Seq may improve therapeutic decision making for most patients with advanced RCC, including patients with non-clear cell RCC for whom no standard treatment is available to data. Prospective clinical trials are needed to evaluate the impact of genomic and transcriptomic diagnostics on survival outcome for advanced RCC patients.

## Background

Renal cell carcinoma (RCC) consists of different histological subtypes (1, 2). The most common histological subtype is clear cell RCC (ccRCC), accounting for approximately 75% of the RCC cases (3). The vast majority of ccRCC is characterized by the loss of the short arm of chromosome 3 (3p)(4), which harbors several tumor suppressor genes. The function of these genes - *VHL, BAP1, PBRM1*, and *SETD2* - is frequently inactivated due to additional somatic mutations or epigenetic changes of these genes on the other allele (4, 5). Although these genetic aberrations can be observed in most patients with ccRCC, the clinical behavior in individual patients differs significantly, from a slowly progressive disease over the years to a rapidly progressive disease with fast clinical deterioration. Therefore, the management of advanced ccRCC varies from active surveillance to systemic therapy.

The therapeutic landscape for patients with advanced ccRCC has changed significantly, in recent years. The introduction of tyrosine kinase inhibitors (TKIs) (6, 7), immune checkpoint inhibitors (ICIs) (8, 9), mammalian target of rapamycin (mTOR) inhibitors (10), and combinations of these anti-cancer therapies (11-13), has significantly improved the outcome for patients with advanced ccRCC. However, there are considerable interindividual differences in outcome, and only a minority of patients experience durable responses (14). For patients with advanced ccRCC, treatment decision making is guided by the International Metastatic RCC Database Consortium (IMDC) criteria (15-17). These criteria include only clinical patient characteristics (i.e. hemoglobin level, time from diagnosis to start of systemic therapy, Karnofsky performance state, calcium level and neutrophil and platelets count).

Moreover, non-clear cell RCC (nccRCC) is a heterogeneous group of different histological subtypes, including papillary and chromophobe RCC (18). As holds for advanced ccRCC, the course of nccRCC differs significantly between patients (19, 20). Since the nccRCC subtypes are considered rare diseases, randomized phase three clinical trials are still lacking for nccRCC (21). As a result, no clear standard of care for patients with advanced nccRCC has been defined (20, 22).

The development of RCC, including its metastatic potential and response to treatment, could largely be explained by the different genomics (2, 4) and evolutionary pathways (5, 23) of this disease. Previous studies have focused on the molecular characterization of primary RCC (2, 4, 24) and the genomic evolution of ccRCC (4, 5, 23). For example, RNA expression analysis in ccRCC has identified different immunogenic and angiogenic gene expression signatures (24, 25). To improve the individualized treatment strategy and survival outcomes for patients with ccRCC and nccRCC, more insight into the genomic make-up of advanced RCC is required.

The objective of this study was to describe the genomic landscape of advanced RCC, by combining whole-genome sequencing (WGS) with matched RNA sequencing (RNA-Seq) data. First, WGS was applied to characterize the genomic make-up of RCC and to identify potential actionable targets for systemic treatment in individual patients with ccRCC and nccRCC. Next, both WGS and matched RNA-Seq data were combined for patients with ccRCC and papillary RCC (pRCC). The RNA-Seq data were applied to cluster RCC based on immunogenic and angiogenic gene expression patterns, aiming to identify those patients who could benefit from either treatment with anti-angiogenic drugs, ICIs or a combination of these therapies and contribute to the next steps in personalized medicine in patients with RCC.

## Methods

### Patient cohort, study procedures, sample collection, clinical data

All patients provided written informed consent for participation in the prospective multicenter Center for Personalized Cancer Treatment (CPCT-02) study (NCT01855477) by the Declaration of Helsinki. The CPCT-02 trial was approved by the medical ethical committee of the University Medical Center Utrecht and the Netherlands Cancer Institute, and local approval was provided for each participating site. Details regarding inclusion criteria, the study protocol, sampling, and sequencing have been previously described (26). In summary, for this analysis, core needle biopsies from the tumor lesion, peripheral whole blood samples and clinical data were collected across 24 hospitals in the Netherlands. Response to treatment was determined according to RECIST v1.1 (27). WGS data from 103 biopsies of 101 patients with advanced RCC were made available. Only one sample per patient was selected for the genomic analyses. After checking the informed consents and the pathological reports, we selected 91 WGS samples (51 previously described by Priestley *et al. (26)*, supplementary data file) and 28 completely new matching RNA-Seq samples.

### Pathological diagnosis

To confirm the histopathological diagnosis of RCC, pathology reports were requested via PALGA, the nationwide network and registry of histo- and cytopathology in the Netherlands (28). Slides and tissue blocks were not available for pathological revision. Alternatively, the pathology reports were reviewed by a genito-urinary pathologist (GvL) to determine whether the microscopic description and immunohistochemistry were compatible with the original diagnosis. The following subtypes were annotated: clear cell, papillary, chromophobe, tubulocystic, and collecting duct carcinoma. Histopathologically confirmed RCC of which the subtype remained unclear was categorized as undefined subtype.

### Whole genome sequencing and preprocessing

Between the 8^th^ of August 2016 and the 3^rd^ of October 2019, tumor and whole-blood pairs were whole-genome sequenced at the Hartwig Medical Foundation (HMF) central sequencing center. When multiple biopsies of one patient were available, the sample covering the most clinical information and/or the sample with the highest estimated tumor cell percentage was selected. A HiSeqX system was applied and 2 × 150 base read pairs were generated using standard settings (Illumina, San Diego, CA, USA). Preprocessing was performed as described by Priestley *et al* (26). Briefly, read pair mapping was performed using BWA-mem (29) to the reference genome GRCh37 (human) with subsequent systematic variant calling and several quality control and/or correction steps. The Genome Rearrangement IDentification Software Suite (GRIDSS) (30) was used for structural variant (SV) calling and LINX (v1.11) (30) for gene fusion event calling. Computational ploidy estimation and copy-number (CN) assessment was performed using the PURPLE (PURity & PLoidy Estimator) pipeline (30), estimating tumor purity and CN profile by combining B-allele frequency (BAF), read depth, and SVs.

### Somatic variant annotation and filtering

As part of the data request, somatic variants were determined using Strelka and provided by the HMF. Variant Call Format (VCF) files with somatic variants were annotated based on GRCh37 with HUGO gene symbols, HGVS notations, gnomAD (31) frequencies using VEP (32) (database release 95, merged cache), with setting “--per_gene”. Exclusively somatic single-nucleotide variants (SNVs), small InDels, multi-nucleotide variants (MNVs) with ≥ 3 alternative read observations and passing variant caller quality control were included in the analyses. Furthermore, population variants were removed to prevent germline leakage, based on the gnomAD database (v2.0.2) (31): gnomAD exome (ALL) allele frequency ≥ 0.001; and gnomAD genome (ALL) ≥ 0.005. Variants specific to the Dutch CPCT cohort were removed based on a panel-of-normals from 1,762 representative normal blood HMF samples. The most deleterious mutation was used to annotate the overlapping gene for each sample.

### Tumor Mutational Burden calculation

The number of mutations per megabase pair was calculated as the amount of somatic genome-wide SNVs, MNVs, and InDels divided by the number of callable nucleotides in the human reference genome (GRCh37) FASTA file:

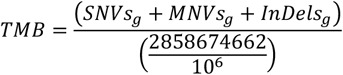

### Ploidy and copy-number analysis

Broad and focal somatic CN alterations in ccRCC were identified by GISTIC2.0 (33) (v2.0.23), using the following parameters: genegistic 1, gcm extreme, maxseg 4000, broad 1, brlen 0.98, conf 0.95, rx 0, cap 3, saveseg 0, armpeel 1, smallmem 0, res 0.01, ta 0.1, td 0.1, savedata 0, savegene 1, qvt 0.1. The distinction between shallow and deep CN events per region was based on thresholding performed by GISTIC2.0. The alterations were assigned a score taking both the amplitude and the frequency of its occurrence across samples into account (G-score). Thresholding was divided into five CN categories, two for deletions (−2 = deep, possibly homozygous loss, -1 = shallow, possibly heterozygous loss), one for diploid (0 = diploid) and two for amplifications (1 = few additional copies, often broad gain, 2 = more copies, often focal gain). Annotation of GISTIC2.0 peaks was performed as follows: A) Wide peaks were annotated with all overlapping canonical UCSC genes within the genomic limits of the said peak. B) Focal peaks were annotated based on overlapping genomic coordinates, using custom R scripts and UCSC gene annotations. As a separate analysis, GISTIC2.0 was executed with ‘brlen’ set to 0.5, calling arm-level events if at least 50% of the chromosomal arm was affected by broad CN events.

### Structural variant analysis

SVs affecting genes were imported using custom R scripts, overlapping genes on at least one breakpoint, using GRCh37 genomic coordinates. SVs with an upstream or downstream Tumor Allele frequency (TAF) below 0.1 as determined by PURPLE and GRIDSS (30) were discarded along with SVs that affected all gene exons. In the case of both (multiple) mutations and/or SVs in the same gene, these were annotated as ‘multiple mutations’.

### Fusion gene analysis

WGS-based LINX TSV files were imported using R and overlapped with the three pillars of ChimerDB (34); deep sequencing data (ChimerSeq), text mining of PubMed publications (ChimerPub), with extensive manual annotations (ChimerKB). Events not present in any pillar of ChimerDB and intra-gene fusions were filtered out. RNA-Seq based fusion genes detected with Isofox (https://github.com/hartwigmedical/hmftools/tree/master/isofox) were imported using R and overlapped with the fusion events detected in the DNA sequencing.

### Recurrent non-coding (and coding) somatic variants

All filtered non-coding somatic variants, occurring in at least four samples were selected as recurrent. Since none of these non-coding recurrent variants were located in regulatory elements (except for the *TERT* promoter) of genes known to be relevant in RCC, we chose to only communicate the *TERT* hotspot. Using the same methodology for coding variants, selecting those arising in at least four samples, did not result in pertinent findings.

### Somatic Driver Genes Analysis

On the ccRCC samples we utilized the dN/dS model (192 Poisson rate parameters; under the full trinucleotide model) to identify genes undergoing mutational selection in ccRCC using the R package dndscv (35) (v0.0.1.0). The substitution and InDel models were used and corrected for sequence composition, gene length and mutational signatures (COSMIC database, see supplementary files). These models test the ratio between nonsynonymous (missense, nonsense and essential splice site) and background (synonymous) mutations. A *q*-value < 0.05 (including and excluding the InDel model) was used to identify genes that drive selection.

### Mutational signatures analysis

Mutational signatures analysis was performed using the MutationalPatterns R package (v3.2.0)(36). The mutational signatures based on single base substitutions (*N* = 90 v3 signatures) were downloaded from COSMIC (37). SNVs were categorized according to their respective trinucleotide context (GRCh37) into a mutational spectrum matrix M_ij_ (where *i* represents 1:96 trinucleotide contexts and *j* represents the number of 1:91 samples). Subsequently, a constrained linear combination of the ninety mutational signatures was constructed per sample using non-negative least squares regression implemented in the R package pracma (v2.2.9). Mutational signatures were bootstrapped (*N* = 100) with MutationalPatterns and argument ‘method’ set to “strict” to assess calling stability. Signature contribution for each sample was determined per 100 samplings, per signature.

### Chromothripsis

Chromothripsis (CT), also known as chromosomal shattering, followed by seemingly random religation, was detected using Shatterseek (38) (v0.4) with default settings. The following definition of CT was employed: (1) ≥25 intrachromosomal SVs involved in the event; (2) ≥7 oscillating CN segments (2 CN states) or ≥14 oscillating CN segments (3 CN states); (3) CT event involving ≥20 Mb; (4) satisfying the test of equal distribution of SV types (*p*-value > 0.05); and (5) satisfying the test of nonrandom SV distribution within the cluster region or chromosome (*p*-value ≤ 0.05).

### Actionable targets

iClusion (https://iclusion.com) data, which connects specific or gene-level aberrations to clinical cancer studies, were provided by HMF. This integrates clinical interpretations from Precision Oncology Knowledge Base (OncoKB) (39), Clinical Interpretation of Variants in Cancer (CIViC) (40) and Cancer Genome Interpreter (CGI) (41). All targets and biomarkers were overlapped with filtered molecular data to verify presence. No RCC-specific effectiveness is taken into consideration with this approach, purely resting the link on the molecular basis of the target to broaden the horizon of actionable targets. Studies included in the databases mentioned can conceivably show associations between drugs and targets in other cancer types than RCC or the subtypes. Targets marked as “gene-level” were generalized for other variation in those genes, not listed in the iClusion data. The identified targets were assessed and manually categorized into the following three categories: on-label drugs for RCC, off-label available, and investigational drugs. Drugs were considered on-label when approval was given for any subtype of RCC in the Netherlands. Whether drugs were on- or off-label available in the Netherlands is defined by the Dutch Medicines Evaluation Board (“College ter Beoordeling van Geneesmiddelen”) (42). This evaluation board considers previous approvals by the U.S. Food and Drug Administration (FDA) and/or European Medicines Agency (EMA).

### Germline analysis

Known pathogenic germline variants (GRCh37) related to cancer and/or Von Hippel-Lindau syndrome were retrieved from ClinVar (43) that were less than 51 bp long, with a review status of “practice guideline”, “expert panel”, “multiple submitters” or “at least one star”. These ClinVar variants were used as filter for the import of germline variants from the VCF files of our cohort. Variants with at least two reads and passing variant caller quality control were included. Furthermore, variants annotated with “high” impact, in genes with known germline variation in RCC (*FH, SDHA, SDHB, SDHC, SDHD, TCEB1, FLCN, CHEK2*) were included.

### RNA sequencing

RNA was isolated from biopsy using the QIAsymphony RNA Kit (Qiagen, Hilden, Germany) for tissue and quantified on the Qubit. Between 50 and 100 ng of RNA was used as input for the KAPA RNA HyperPrep Kit with RiboErase (Human/Mouse/Rat) library preparation (Roche) on an automated liquid handling platform (Beckman Coulter). RNA was fragmented (high temperature in the presence of magnesium) to a target length of 300 bp. Barcoded libraries were sequenced as pools on either a NextSeq 500 (V2.5 reagents) generating 2 × 75 base read pairs or on a NovaSeq 6000 generating 2 × 150 base read pairs using standard settings (Illumina, San Diego, CA, USA). BCL output from the sequencing platform was converted to FASTQ using Illumina’s bcl2fastq tool (versions 2.17 to 2.20) using default parameters. RNA-Seq data were aligned using STAR (44) to GRCh37 resulting in unsorted BAMs including chimeric reads as output. Gene and transcript counts were generated and used for subsequent fusion detection using Isofox (https://github.com/hartwigmedical/hmftools/tree/master/isofox).

### RNA sequencing analyses

Raw read counts were imported in R and filtered on protein coding genes based on Ensembl GTF file (45) (Homo sapiens GRCh37, version 87). *t*-distributed stochastic neighbor embedding (*t*-SNE) was performed on variance stabilized read counts (generated by DESeq2 varianceStabilizingTransformation) of all protein coding genes. Differential expression analysis between ccRCC and pRCC was performed on raw read counts using DESeq2 (46) with the Wald-test. Statistical significant results with Benjamini-Hochberg adjusted *p*-value < 0.05 were further filtered to base mean > 100 counts and absolute log_2_ fold change ≥ 1. The heatmap with the top most significantly differentially expressed genes (based on the lowest adjusted *p*-value) was made using variance stabilized read counts and Euclidean distances on scaled data. Gene signature heatmap was produced with centered Z-Scores with Euclidean distances. Gene set enrichment analyses were performed using fgsea (47) (Monte Carlo approach with Adaptive Multilevel Splitting) with MSigDB (48) Hallmarks and Reactome pathways (49) as gene sets. Reproduction of the D’Costa *et al*. gene signature (25) was done using 65 of 66 original genes, since *PECAM1* was on a genome patch not included in the RNA-Seq mapping supplied by HMF. Heatmaps were produced using pheatmap with Ward.D clustering.

### Data and material availability

Data were provided by HMF, which were used under data request number DR-088 for the current study. Both WGS, RNA-Seq and clinical data are freely available for academic use from the HMF through standardized procedures and request forms can be found at https://www.hartwigmedicalfoundation.nl. All tools and scripts used for processing the WGS data are available at https://github.com/hartwigmedical/ and/or can be provided by authors upon request.

## Results

### Patient selection

In total, WGS data of 91 patients with histopathologically confirmed RCC from 24 hospitals in the Netherlands were included in the analyses (**Figure 1A**). Matched RNA-Seq data were available for 28 patients (**Figure 1A**). Overall, 72 patients were diagnosed with ccRCC, nine patients with pRCC, one with chromophobe RCC, one with tubulocystic RCC, and one with collecting duct carcinoma. The RCC subtypes of the remaining seven patients could not be further defined pathologically. The main biopsy sites were the kidney (*N* = 24), bone (*N* = 15), and lymph nodes (*N* = 14) (**Figure 1B**). The median age of patients at time of the biopsy was 65 years (range 40-83), 79% of the patients were male, and 78% of the patients did not receive any systemic treatment before biopsy (**Table 1**). Most patients (84/91, 92%) were treated with systemic therapy after the biopsy was collected. This treatment consisted mostly of TKIs (61/84, 73%) or ICIs (19/84, 23%), whereas the remaining patients received combination treatment (4/84, 5%).

**Figure 1.**
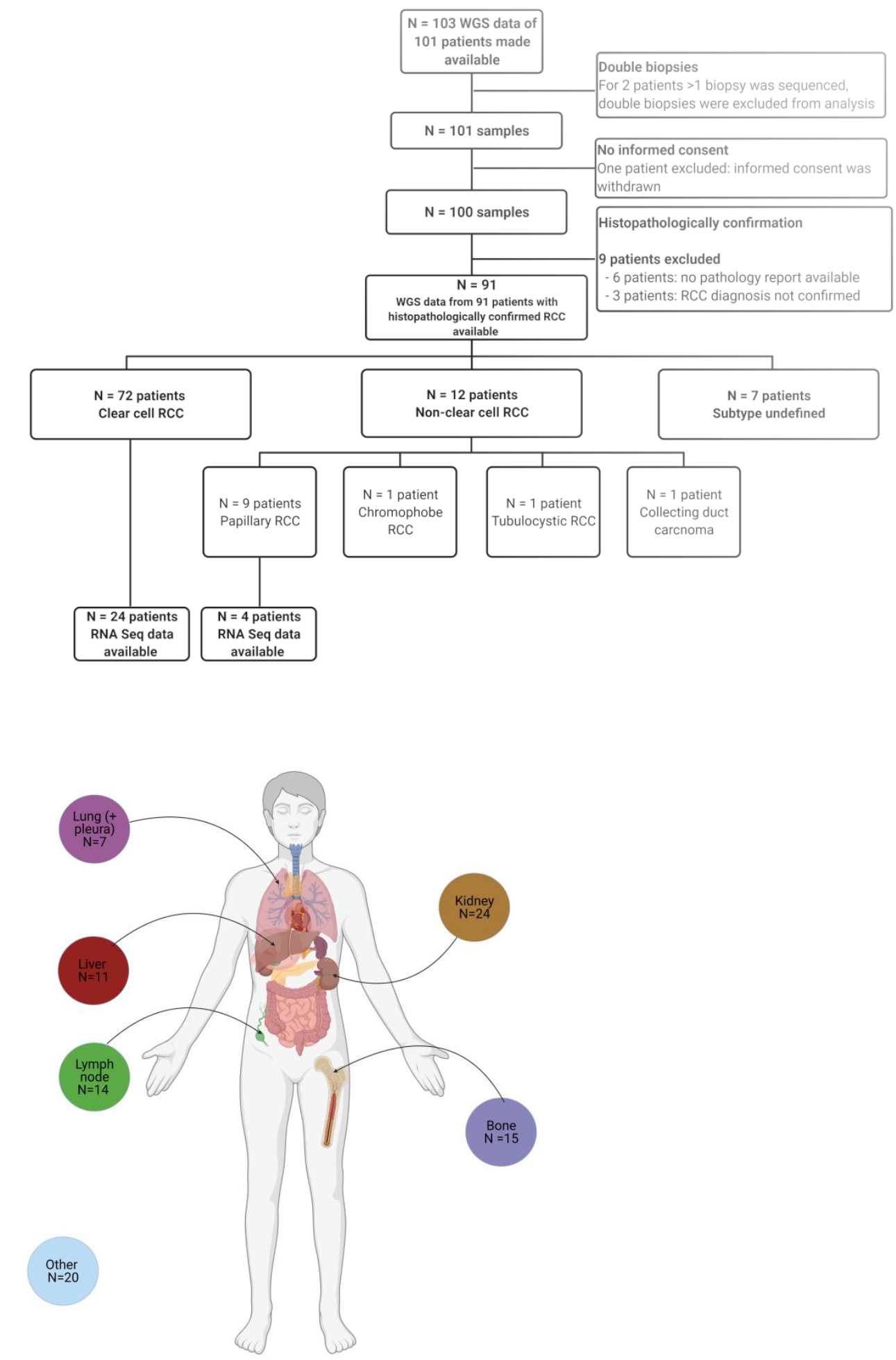
Overview of sample selection and biopsy sites. **1A** illustrates the selection of samples for the WGS (*N* = 91) and matched RNA-Seq (*N* = 28) analyses. In **1B**, the main biopsy sites are shown. Next to the illustrated biopsy sites, other biopsy sites (*N* = 20) include biopsies from soft tissue, muscle biopsies and biopsies from subcutaneous tissue. Figure **1B** was created with biorender.

**Table 1.** Overview of patients’ characteristics.

|  | Frequency | Percentage |
| --- | --- | --- |
| **Sex** |  |  |
| Male | 72 | 79% |
| Female | 19 | 21% |
| **Median age 65 years [range 40-83]** | ----- | ----- |
| **Histological subtype** |  |  |
| Clear cell RCC (ccRCC) | 72 | 79% |
| Non-clear cell RCC (nccRCC) | 12 | 13% |
| Papillary RCC (pRCC) | 9 | 75% |
| Chromophobe RCC (chRCC) | 1 | 8% |
| Collecting duct carcinoma (CDC) | 1 | 8% |
| Tubulocystic RCC (tRCC) | 1 | 8% |
| Undefined subtype | 7 | 8% |
| **Prior systemic treatment (n=number of lines)** |  |  |
| No | 71 | 78% |
| Yes (1) | 10 | 11% |
| Yes (≥2) | 10 | 11% |
| **Treatment after biopsy (N = 84)** |  |  |
| **Tyrosine kinase inhibitors (TKIs)** | 61 | 73% |
| Pazopanib | 36 | 59% |
| Sunitinib | 23 | 38% |
| Cabozantinib | 1 | 1.6% |
| Lenvatinib | 1 | 1.6% |
| **Immune checkpoint inhibitors (ICIs)** | 19 | 23% |
| Nivolumab monotherapy | 13 | 68% |
| Nivolumab + ipilimumab | 6 | 32% |
| **Combination treatment** | 4 | 5% |
| Avelumab + axitinib | 4 | 100% |

### Whole-genome sequencing analyses

The tumor and blood biopsies were whole genome sequenced (WGS) with a median coverage of 102X and 38X (Supplementary figure 1A). The number and type of somatic changes in the WGS data were analyzed on both small mutations and structural variations (supplementary figure 2), showing similar numbers as previous RCC cohorts (7,224 substitutions, Q1–Q3: [5648-8581] *vs*. 7,050, Q1–Q3: [6434-9504], *p-*value = 0.45, Mann-Whitney U test (5)).

Somatic small variants, copy-number alterations (CNAs) and structural variants (SVs) were identified as described previously (26). The median tumor mutational burden (TMB) of patients with ccRCC was 2.8 [interquartile range (IQR) 1.1] (**Figure 2A**). Only two patients (one with ccRCC and one with an undefined subtype of RCC) had a TMB > 10 bases. In the two samples with high TMB, mutational signatures were related to defective DNA mismatch repair, covering more than 25% of their single-nucleotide variants (SNVs) (**Figure 2A/E**). Interestingly, the lowest TMB (i.e. 0.16) was detected in a sample from a patient with tubulocystic RCC (*N* = 1) which had minimal genomic aberrations in general, and only two detectable SVs (**Figure 2C/D;** tandem duplication and break-end). SBS40 (unknown etiology) was the dominant mutational signature, with a mean relative contribution of 74% in this cohort. To assess the reliability of the SBS40 calling, the mutational signature calling was bootstrapped, which revealed a very high variance (median difference in the assignment of 37%) in the relative contribution of SBS40 (Supplementary figure 3).

**Figure 2.**
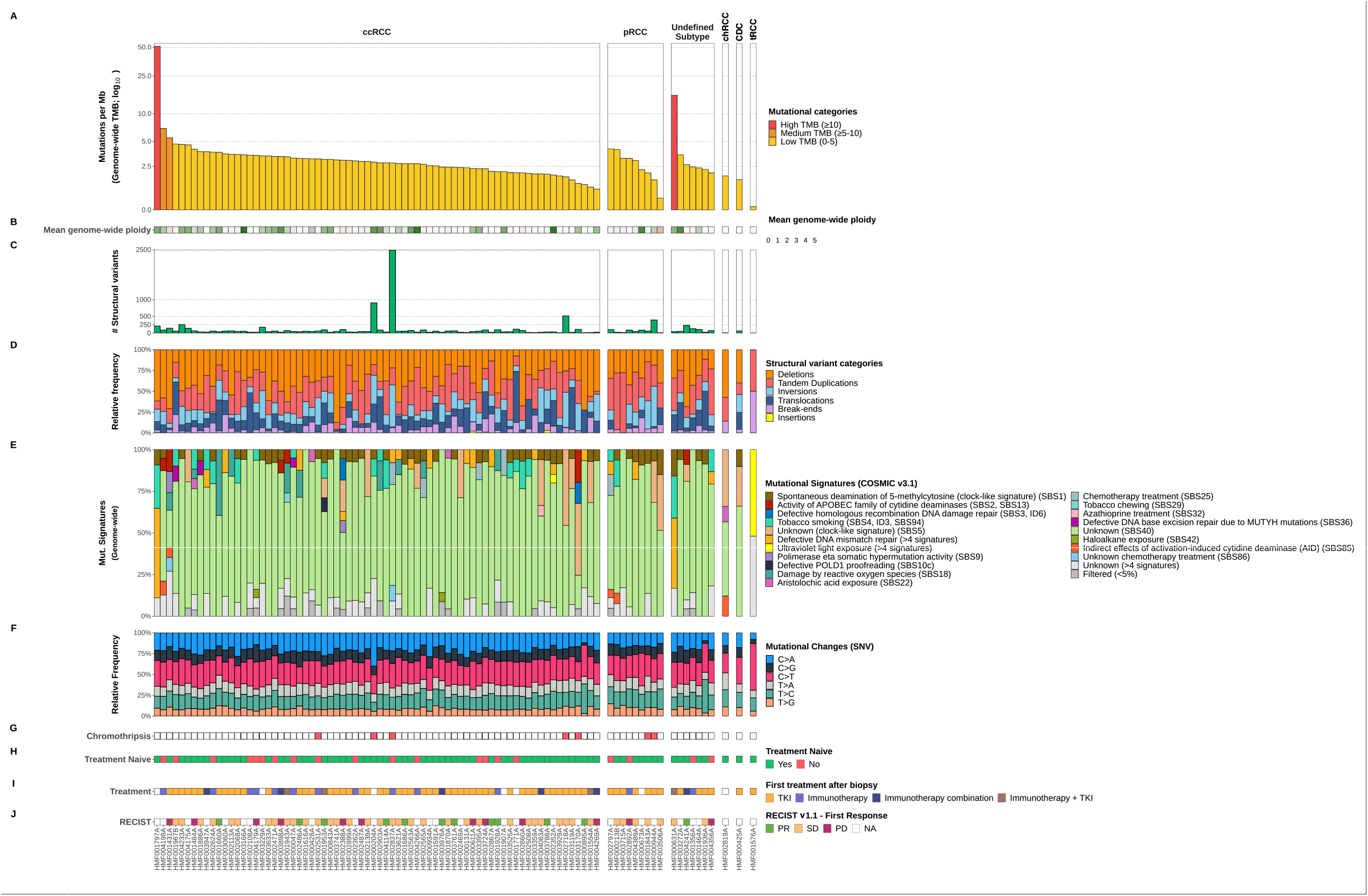
Overview of genomic characteristics of whole-genome sequenced advanced RCC cohort (*N* = 91) Track **A** shows the tumor mutational burden (mutations per Mb; yellow for low (0-5), orange for medium (5-10) and red for high (> 10)). Track **B** shows the mean genome-wide ploidy, with white representing diploidy. Tracks **C** and **D** illustrate the abundance of structural variants and the relative frequency of the types of these variants. Tracks **E** and **F** show the relative mutational signature contribution (COSMIC signatures v3) and the relative frequency of mutational changes at the base level. Track **G** shows the presence of chromothripsis. Track **H** shows whether patients were treatment naive at the time of biopsy. Tracks **I** and **J** indicate the first treatment given after biopsy (if any) and the first tumor response according to RECIST v1.1, respectively. On the x-axis, the figure is arranged in descending order by tumor mutational burden per RCC subtype. ccRCC = clear cell renal cell carcinoma. pRCC = papillary renal cell carcinoma. Undefined subtype = renal cell carcinoma, with undefined subtype. chRCC = chromophobe renal cell carcinoma. CDC = collecting duct carcinoma. tRCC = tubulocystic renal cell carcinoma. NA = not available.

The frequency of genomic SNVs, multi-nucleotide variants (MNVs), InDels, and the collective coding mutations showed a similar pattern across the different RCC subtypes (Supplementary figure 2, supplementary data file). In total, 713,077 somatically acquired SNVs, 173,579 InDels and 9,964 MNVs were detected in the RCC genomes. Transversions were more frequently found than transitions in ccRCC, pRCC and undefined subtypes (Supplementary figure 2B and 2E). Missense variants were the most dominant protein variant type and accounted for > 60% of the small variants in all subtypes (Supplementary figure 2G). For 57% of ccRCC, the genome-wide ploidy was 2, while 32% had a genome doubling with a ploidy of 3 or higher (Supplementary figure 2D). Differences in DNA ploidy have been associated with tumor differentiation and diploidy has been related to well-differentiated RCC (50). Considering SVs, a total of 3,121 deletions, 941 translocations, 2,196 tandem duplications, four insertions, 2,714 inversions and 641 break-ends were detected (Supplementary Fig 2F). The number of patients with events considered chromothripsis was limited and present in only five patients with ccRCC, two patients with pRCC and absent in the other subtypes. Furthermore, these chromothripsis events primarily did not involve the classic t(3;5) event. Only one of the samples with chromothripsis did have a t(3;5) translocation (Supplementary figure 4), while these specific translocations were more frequently detected in the cohort of patients without chromothripsis events (29.7%, Supplementary figure 5). While clinical data concerning pre-treatment (**Figure 2H**), first treatment after biopsy (**Figure 2I**), and accompanying RECIST score (**Figure 2J**) was available, no correlative conclusions were drawn due to heterogeneity which would undermine any reliability attached to the data (Supplementary figure 6).

In patients with ccRCC, both previously described and novel amplifications and deletions were detected. Statistically significant CNA peaks (A) and arm-level copy-number alterations (B) in ccRCC are presented in Supplementary figure 7. Arm-level CNAs covering more than 50% were detected for amplifications of 1q, 5q, 7q, 8q, 12p and 20q, and deletions in 3p, 9p and 14q. All these CNAs have been described previously(51). Furthermore, in the current cohort, amplifications of 5p, 7p, 12q, 16p and 20p, and deletions of 4p, 4q, 6p, 8p, 9q, 14p, 18p and 18q were also statistically significant.

Next, the WGS data were analyzed on driver genes of the ccRCC samples using the dN/dS algorithm and on CNAs by GISTIC 2.0. The driver gene analyses revealed that most driver genes in ccRCC encompassed variations in the chromosome 3p region: *PBRM1* (94.4%), *VHL* (93.1%), *SETD2* (90.3%), and *BAP1* (87.5%) together with events in tumor suppressor genes on other chromosomes, such as *CDKN2A*/*B* (66.7%) and *PTEN* (30.6%) (**Figure 3**, supplementary data file). In addition, many patients with ccRCC had focal deletions in genes described as putative tumor suppressor genes, e.g. *PTPRD* (65.3%) and *NEGR1* (44.4%) (4, 52). Deletions of *PTPRD* have been described as a potential risk factor for the development of ccRCC (52, 53). Moreover, amplifications were present in genes associated with cell proliferation and angiogenesis, such as *CDK6* (55.6%, also prevalent in pRCC at 77.8%) and *CCNE1* (16.7%) (54, 55). Pathogenic germline mutations related to cancer or Von Hippel-Lindau syndrome were found in nine patients in different RCC subtypes and included *ATM, FLCN, CHEK2, FH, SDHA* and *MITF* (**Figure 3**).

**Figure 3.**
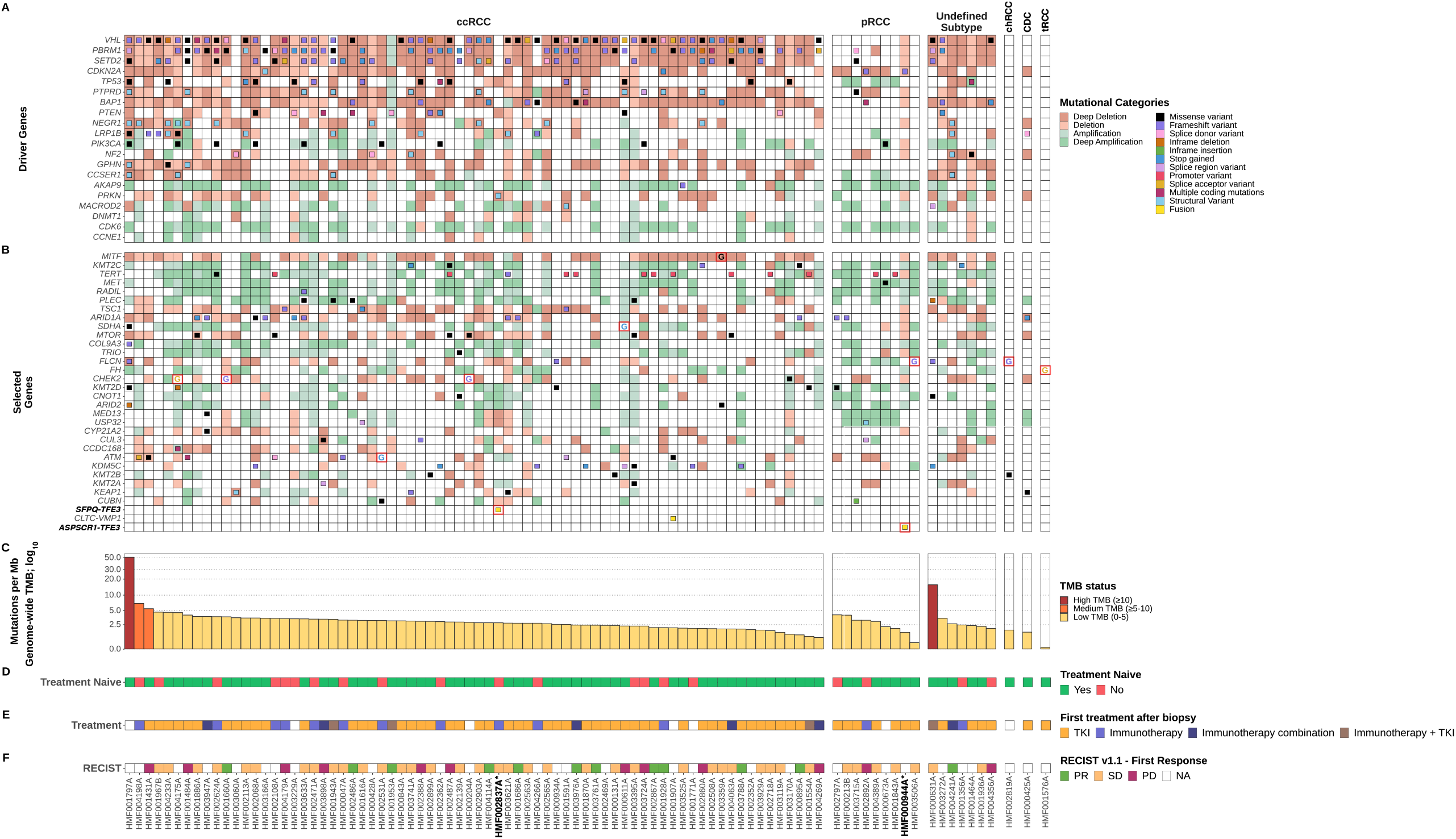
Overview of coding mutations and copy-number alterations in driver genes in whole-genome sequenced advanced renal cell carcinoma cohort (*N* = 91) The oncoplot in track **A** shows mutations (filled center) and copy-number alterations (grid cell background) of driver genes determined by dN/dS and GISTIC2.0. Track **B** also shows an oncoplot, but on selected genes, not passing any statistical threshold. Germline pathogenic mutations are indicated with the capital letter ‘G’ (and red border), utilizing the same color coding as the somatic mutations. Consequential fusion genes are indicated in yellow, with a red border. Track **C** shows the tumor mutational burden (mutations per Mb; yellow for low (0-5), orange for medium (5-10) and red for high (> 10)). Tracks **D, E** and **F** shows whether patients were treatment naive at time of biopsy, if systemic treatment was given after time of biopsy, and the first tumor response after systemic treatment according to RECIST v1.1, respectively. Bold sample names with asterisks indicate MiT family translocation RCC. The figure is arranged in descending order by tumor mutational burden per RCC subtype on the x-axis.

In addition, genes that were previously described as frequently mutated somatically (*q*-value < 0.05) in (cc)RCC by Braun *et al*. (56) and in pRCC by Turajlic *et al*. (57), that were not statistically significant in our driver gene analysis, were included to extend our analysis. The commonly most affected and added genes in ccRCC were *MITF* (3p), *TERT* (5p), *RADIL* (7p), and *MET* (7q), predominantly as bystanders of arm-level events. The *TERT* promoter hotspot variant (C228T) (5) was detected in both ccRCC (*N* = 10) and pRCC (*N =* 2). *MET* and *RADIL* were also frequently affected in pRCC (both in 77.8%), along with *MED13* and *USP32* (both in 66.7%). Genes associated with poor prognosis in ccRCC showed a low mutational frequency in our cohort (4), such as the Krebs cycle genes (e.g. *SDHA, FH*).

Furthermore, previously validated fusion events (34) were detected, with *CLTC-VMP1* and *SFPQ-TFE3* both occurring once in the ccRCC group, along with a fusion event of *ASPSCR1-TFE3* in one patient with pRCC. The histopathological diagnosis had to be reconsidered for both patients with a detected *TFE3* fusion. As a result, these patients were reallocated in a different subcategory of RCC, i.e. MiT family translocation renal cell carcinomas (1). Overall, characteristic clear cell driver gene events — such as somatic *VHL* mutations/deletions — were infrequent in nccRCC, except for *CDKN2A* deletions in pRCC.

Next, we investigated if the tumors contained targetable variants (**Figure 4**), that are known drivers in RCC (**Figure 3**) but also proteins which are considered targetable in other cancer indications. For example, specific drugs have been developed for target-specific variants encoded by *CDK4/6* or *EGFR* (58-60). This may result in off-label availability of these drugs for patients with similar aberrations in RCC, e.g. in the context of a clinical trial (61). Furthermore, somatic aberrations in cancer genes, e.g. *TP53*, are also considered biomarkers for targeted treatments (62). Lastly, some variants leading to specific mechanisms were also considered targetable for treatment. For instance, TKIs targeting VEGF signaling are known to be effective in patients with *VHL* mutations and are on-label available for patients with RCC (63). Overall, for the majority (90 out of 91) of patients in this cohort actionable targets were detected, even for patients with nccRCC for whom no standard treatment is available to date.

**Figure 4.**
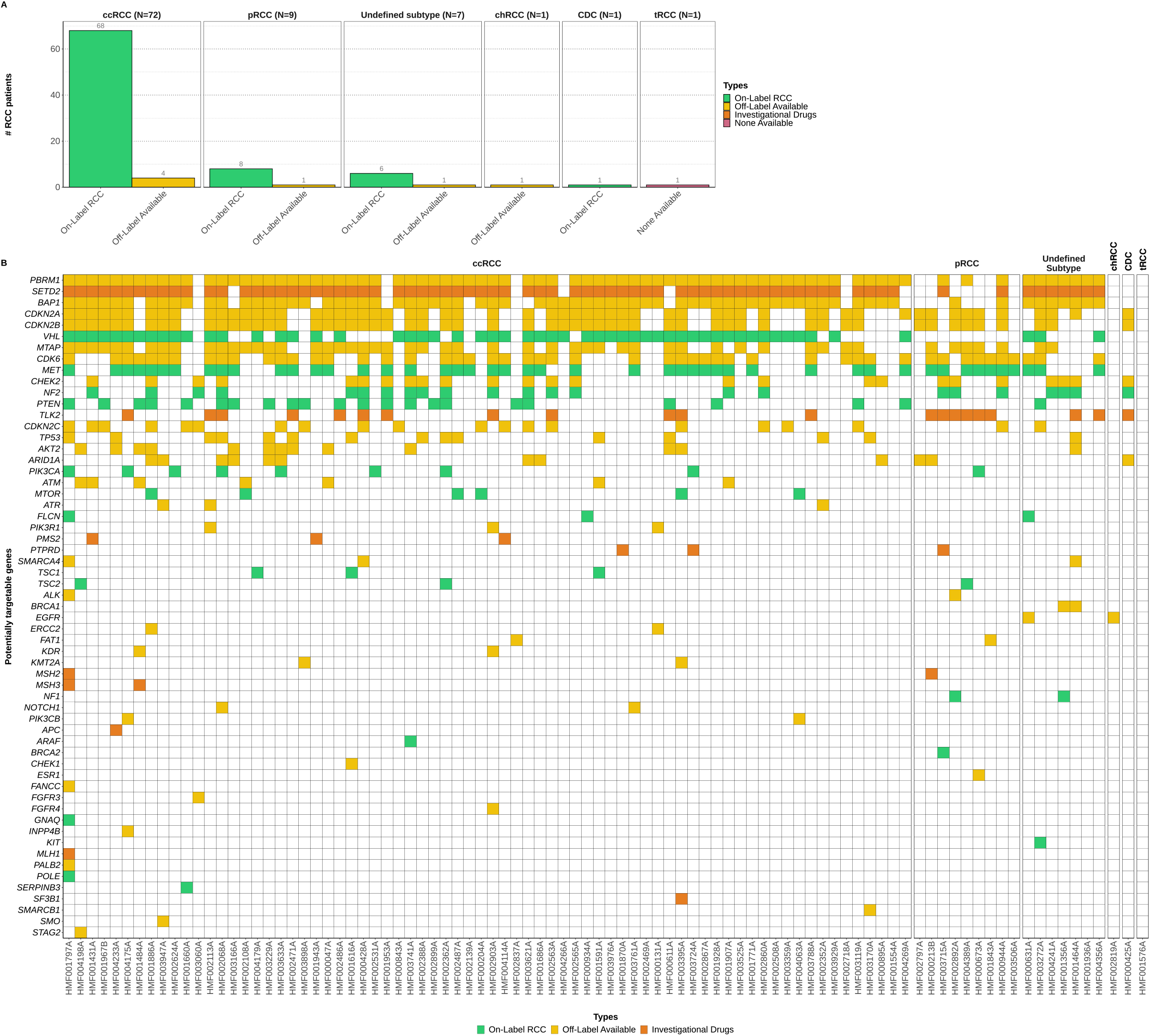
Overview of DNA-based biomarkers and potential treatment options in the whole-genome sequenced advanced renal cell carcinoma cohort (*N* = 91) Track **A** Percentage of potentially available treatment options based on genomic characteristics. Treatment options are categorized according to the highest level of drug availability in clinical practice (on label available – off label available – investigational drugs). Track **B** Potentially actionable alterations at gene-level with each column representing a sample, ordered descendingly by tumor mutational burden per subtype on the x-axis. A detailed description of actionable targets can be found in Supplementary Data file 1.

### Transcriptome analyses of advanced RCC

Differential Expression Analysis (DEA) was performed on RNA-Seq data (Supplementary figure 1B) to discriminate the two most frequently diagnosed histological subtypes, ccRCC (*N* = 24) and pRCC (*N* = 4). Next to the *t*-distributed stochastic neighbor embedding (*t*-SNE), which showed a clear separation between the two subtypes (Supplementary figure 8), the DEA resulted in 1,546 significantly (adjusted *p*-value < 0.05) differentially expressed genes. The hundred genes with the smallest adjusted *p*-value are shown in **Figure 5A**. In this top hundred list, several genes are known to be associated with the development or course of RCC. For instance, *LOX* (64) and *MAPKAPK3* (65) correlate with poor survival in RCC. In addition, various other genes were differentially expressed and have been described in other malignancies, such as *TUSC2* (66), *CAPN1* (67), *PCSK6* (68), and *CD2* (69). The differential expression of these genes confirmed a clear distinction between pRCC and ccRCC at the transcriptomic level.

**Figure 5.**
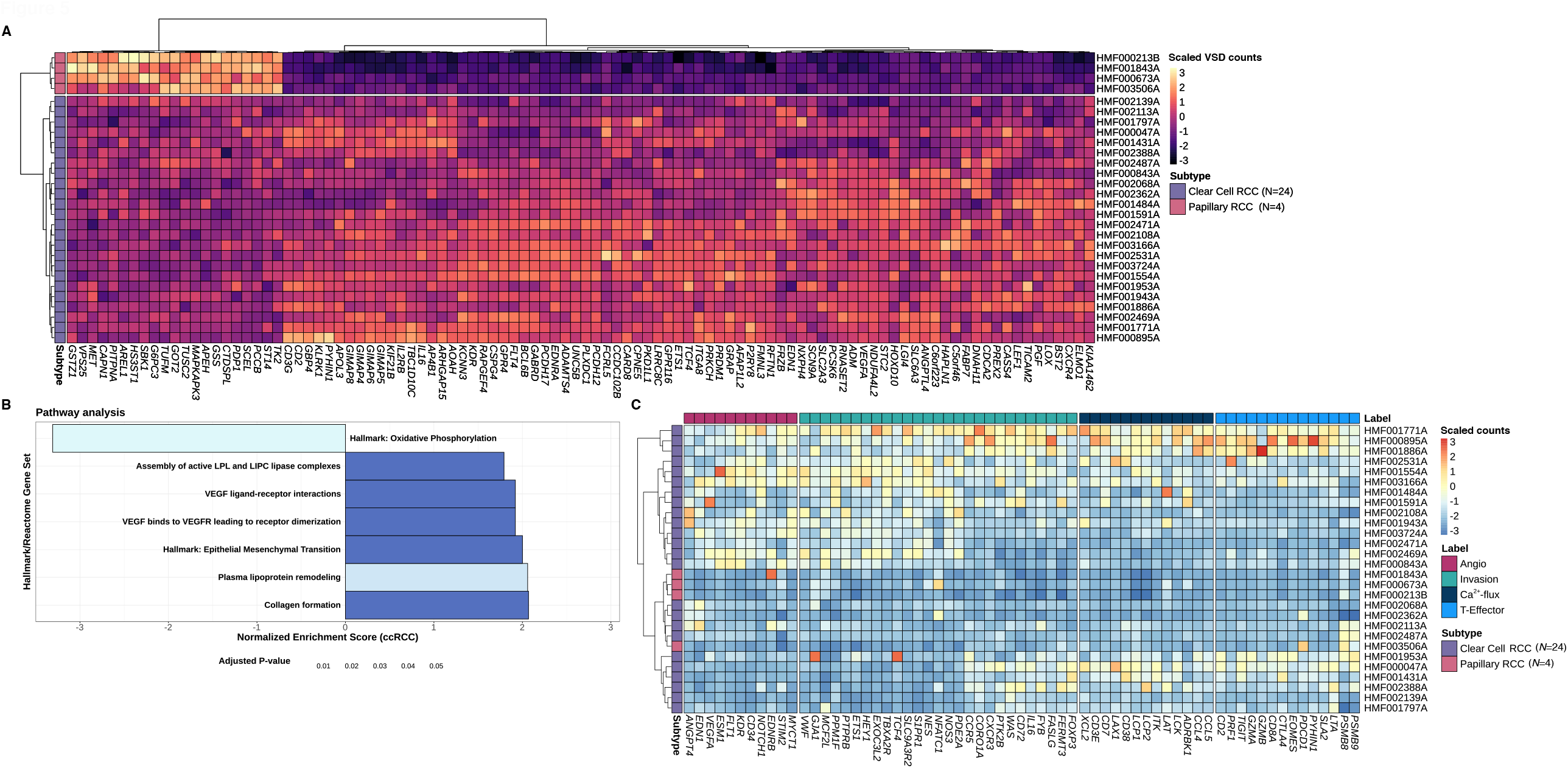
RNA sequencing cohort and differential expression analysis between clear cell renal cell carcinoma (ccRCC) and papillary RCC (pRCC) with classification according to gene signatures^25^. Track **A** shows a heatmap of Z-scores of variance stabilized values with unsupervised clustering of the top 100 transcripts based on the smallest adjusted *p*-value and colored according to Z-scores. Tracks **B** shows gene set enrichments based on sets (y-axis) from the molecular signatures database hallmarks and Reactome pathways, with the normalized enrichment score (NES) on the x-axis. Bar charts are visualized with ccRCC taken as reference (*N* = 24) (positive NES equals expression up in ccRCC and down in pRCC (*N* = 4)). Track **C** shows unsupervised clustering on the rows (patients) with color coding indicating the RCC subtype (purple for ccRCC and pink for pRCC) and colored according to Z-scores. The x-axis has been cut into several gene groups related to angiogenesis, invasion, Ca^2+-^flux and T-effector cells, as defined by D’Costa *et al*. ^25^ and their stated order.

Second, the pathway analysis (**Figure 5B**) of the 1,546 significant differentially expressed genes revealed cancer hallmarks such as oxidative phosphorylation and epithelial-mesenchymal transition (EMT), among others (70). Significant Reactome pathways (49) of differentially expressed genes in ccRCC compared to pRCC samples mainly showed pathways related to VEGF and collagen formation. Pathways related to poor prognosis (4), such as the AMPK complex, the Krebs cycle genes, the pentose phosphate pathway and fatty acid synthesis, were not differentially expressed between ccRCC and pRCC. A heatmap of the top differentially expressed genes between ccRCC and pRCC and a *t*-SNE plot show that RCC samples of undefined subtype cluster with either ccRCC or pRCC samples based on the differential gene expression of these subtypes (Supplementary figure 9).

The 66 gene-signature has been based on previous data from the IMmotion150 trial (25). High expression of ‘angiogenic’ genes and specific ‘invasion’ genes is applied to sub-classify RCC as ‘angiogenic’, which would be predictive for response to TKIs targeting VEGF signaling. In case of high expression in ‘Ca^2+^-Flux’, ‘T-Effector’, and other ‘invasion’ genes, RCC is sub-classified as ‘immunogenic’, indicating a likely response to ICIs. According to this gene signature, ccRCC cases in the current cohort could be classified as either immunogenic or angiogenic (**Figure 5C**). Moreover, all patients with pRCC had low expression of these genes, except for one individual patient with expression of *EDNRB (71)*.

## Discussion

In this study, the genomic and transcriptomic landscape of advanced RCC was characterized for 91 individual patients. First, genomic data showed that, next to *VHL* alterations (93.1%), most common driver gene mutations in ccRCC included alterations in tumor suppressor genes of different pathways such as *SETD2* (90.3%) and *PTEN* (30.6%). While TMB was comparable amongst the different subtypes of RCC, the driver gene analyses showed a distinctive pattern between patients with ccRCC and nccRCC. Furthermore, WGS revealed potential actionable targets for 90 out of 91 patients and WGS might therefore contribute to a more individualized treatment strategy for patients with advanced RCC.

For a subgroup of patients (*N* = 28), transcriptomic data were also generated. RNA-Seq could be applied to distinguish ccRCC, pRCC and histologically undefined RCC based on the differential gene expression. Using the 66-gene signature (25) on the RNA-Seq data, made it possible to sub-categorize ccRCC into immunogenic or angiogenic signatures, whereas classification in pRCC (*N* = 4) using these signatures was not feasible.

At the genomic level, the findings for ccRCC mainly corresponded with previous findings (4, 5, 72). Previously described statistically significant arm-level events were also found in this cohort, i.e., amplifications of 1q, 5q, 7q, 8q, 12p and 20q, and deletions in 3p, 9p and 14q. However, a striking difference in the frequency of some events is caused by a distinct definition of ‘arm-level’. For instance, both full loss of 3p and focal loss of events along 3p21-p25 (*VHL, PBRM1, BAP1, SETD2*) are counted as ‘arm-level’ in the ccRCC characterization from the Cancer Genome Atlas Research Network (4). In contrast, we defined these (and other large) events as covering >50% of the chromosome arm (Supplementary Figure 7). The massive contribution of SBS40 to the mutational landscape of nearly all RCC subtypes in this cohort is remarkable. However, bootstrapping showed that SBS40 was the least robust signature, indicating that this signature could act as a sink for mutations that are difficult to fit. Since CPCT-02 cohorts with other tumor types did not show the high contribution of SBS40 (73, 74), this is certainly not a result of a bias in the sequencing or our workflows. In contrast to other ccRCC studies, the number of chromothriptic events was limited in our study and occurred in only five patients. In previous studies, chromothripsis was defined as the combination of a chromothripsis event and a translocation event with concurrent 3p loss and 5q gain, which were called “t(3;5) chromothripsis events” (5). Although both chromothripsis and translocation (t(3;5)) events occurred in our cohort, for most of the cases, these events were independent of each other (χ^2^ test, p-value = 0.6193, events visualized in Supplementary figures 4 and 5).

At the transcriptomic level, assigning ccRCC biopsies to either immunogenic or angiogenic signatures may indicate which treatment could be most beneficial for individual patients. The introduction of ICIs has significantly changed the therapeutic landscape for patients with advanced ccRCC (75, 76), resulting in a clinical need to select patients who will benefit from angiogenic or immunogenic treatment (17). RNA-seq data could assist clinical decision making when choosing the optimal treatment strategy for the individual patient with advanced ccRCC. For patients with high expression of genes annotated as immunogenic, first–line treatment with ICIs should be considered, whereas, for patients with high expression in angiogenic genes, treatment with a TKI should be taken into consideration. For those patients with low expression throughout all these genes, combination treatment with TKI/ICI may be considered. However, treatment based on actionable targets identified by WGS could be the most effective option. Treatment selection based on gene expression has already shown promising results for patients with RCC (12, 77). However, further research in a prospective setting is still warranted (78).

The distinctive mutational gene pattern between ccRCC and nccRCC showed that these tumors are different entities, while differences within the nccRCC subtypes were also evident. For example, none of the patients with nccRCC had somatic *VHL* alterations and other ccRCC driver genes hardly showed mutations in nccRCC. This is of great importance, as the development of targeted drugs is based on driver mutations or their downstream consequences. For instance, the frequently mutated *VHL* gene in ccRCC has been the basis for the developing angiogenesis inhibitors. The loss-of-function mutations in this tumor-suppressor gene result in the accumulation of *HIF1α/2α*, eventually leading to overexpression of *VEGF/PDGF, AXL*, and *MET*, among others (63, 79). Several TKIs that have been approved for the treatment of advanced ccRCC (6, 7, 63, 80) and interfere at different levels in this cascade (63, 79). More recently, Hypoxia Inducible Factor (HIF) inhibition has also shown proven efficacy in patients with *VHL* alterations (81). However, as patients with nccRCC in our cohort showed no mutations in *VHL* or other angiogenesis related pathways, it is questionable whether treatment directed against the *VHL* pathway would be the most effective therapy for this group of patients. On the other hand, germline mutations associated with specific genetic syndromes were detected in several patients with nccRCC. For instance, the patient with tRCC had a germline *FH* mutation, which makes it likely that this patient suffered from hereditary leiomyomatosis renal cell carcinoma syndrome (HLRCC) (82). Moreover, patients with a germline *FLCN* mutation may suffer from the Birt-Hogg-Dubé syndrome, often associated with chromophobe renal cell carcinoma or oncocytoma (83). Our germline analysis shows that the number of germline mutation is relatively high in the more rare subtypes of RCC, demonstrating the importance of germline mutation analysis for (rare forms of) nccRCC. Since germline mutations play an important role in the development of RCC and are often associated with particular types of RCC, detecting these mutations allows for personalized and targeted treatment of the affected patients.

In clinical practice, RCC is usually defined histopathologically. As a result, there is a large dependency on experienced pathologists. However, discrepancies among these experts remain (84, 85). In our cohort, nearly 8% of RCC cases could not be sub-classified through histopathological assessment. Thereby, for two patients, a fusion gene was predicted using WGS which led to the revision of the original histological diagnosis and allocation to a different subgroup, i.e. MiT family translocation renal cell carcinoma (1). As the histopathological classification defines the treatment strategy, this could have a significant clinical impact. Since different subtypes and growth patterns of RCC are driven by gene expression (86), a next generation sequencing-based classifier could be feasible. Here, we showed that analyses of driver mutations (*VHL, PBRM1, SETD2)* and RNA-Seq data reveal clear differences among the different RCC subtypes. As shown in Supplementary Figures 8 and 9, clustering of the undefined RCC subtypes is feasible and could be useful to clarify the histological subtype in clinical practice.

This study has some important limitations. First, the collected clinical data within the CPCT-02 study were limited and therefore it was not possible to reliably correlate genomic and transcriptomic findings to clinical data. A correlation with clinical data and outcome could have confirmed whether patients with certain gene signatures indeed had benefit from a specific treatment. Therefore, validation in a prospective trial is needed before clinical implementation. Second, the limited number of patients with nccRCC made it challenging to run separate analyses for this group. Since the subgroup of patients with nccRCC consists of less common and heterogeneous subtypes, very little is known about the genomics of nccRCC. Therefore, we decided to include all patients with nccRCC, even subgroups containing only a single patient. Finally, the collected data were heterogeneous, with a broad spectrum of pre-treatment schedule, different treatment strategies after biopsy (**Supplementary figure 6**). All patients had metastatic or locally advanced disease but biopsies taken from metastatic and the kidney (primary site) were both included in the analyses. This strategy could have significantly impacted the analysis. However, previous studies have shown clear consistencies between primary tumor biopsies and their metastasis counterpart ^5^. In addition, the heterogeneity in this cohort reflects the daily clinical practice of patients who present with advanced RCC, including primary metastatic disease. Despite this clinical heterogeneity, a clear genomic and transcriptomic signal could be extracted, indicating that the genomic and transcriptomic analyses are feasible for clinical implementation.

## Conclusions

In conclusion, there are evident genomic and transcriptomic differences between RCC subtypes. The analysis of driver mutations, in combination with the clustering of RNA-Seq data, could assist the histopathological subtyping of RCCs in clinical practice. In addition, RNA-Seq data could identify patients with ccRCC who may benefit more from treatment with either ICIs, TKIs or a combination of these drugs. Genomic and transcriptomic analyses are promising to identify actionable targets and individualize treatment strategies in most patients with RCC, even for patients with nccRCC. Although these results are promising, prospective clinical trials are still needed to evaluate whether genomic and transcriptomic diagnostics contribute to improved survival outcomes in individual patients with advanced RCC.

## Declarations

### Ethics approval and consent to participate

All patients provided written informed consent for participation in the prospective multicenter Center for Personalized Cancer Treatment (CPCT-02) study (NCT01855477). The CPCT-02 trial was approved by the medical ethical committee of the University Medical Center Utrecht and the Netherlands Cancer Institute. Local approval was provided for each participating site.

### Consent for publication

Not applicable.

### Competing interests

K.J. declares travel expenses from Ipsen, outside the submitted work; P.H. declares consultancy roles for Astellas, MSD, Ipsen, Pfizer, AstraZeneca, and Bristol-Myers Squibb, all outside the submitted work; H.M.W. declares honoraria from Roche and Astellas and travel expenses from Ipsen and Astellas, all outside the submitted work; S.F.O declares research grants from Novartis, Pfizer and Celldex Therapeutics and advisory board for Bristol Myers Squibb (all paid to the institution); M.L. declares speakers fee of BMS and advisory board of MSD, both paid to institution; R.H.J.M. declares speakers fee from Novartis and advisory role for Servier, patency from Pamgene, and investigator-initiated research (paid to institution) from Astellas, Bayer, Boehringer-Ingelheim, Cristal Therapeutics, Pamgene, Pfizer, Novartis, Roche, Sanofi, Servier, all outside the submitted work; M.P.L. declares advisory board for Amgen, Astellas, Astra Zeneca, Bayer, INCa, Janssen Cilag BV, MSD, Novartis, Pfizer, Roche, Sanofi, Servier, consulting role for Julius Clinical and Research Grants (paid to institution) from Astellas, Janssen, MSD, Sanofi, all outside the submitted work; H.J.G.W. declares speakers honoraria from Bayer, Depositary receipts for shares from Cergentis B.V. all outside the submitted work; A.A.M.V. reports advisory board (all paid to institution) of BMS, MSD, Merck, Pfizer, Ipsen, Eisai, Pierre Fabre, Roche, Novartis, Sanofi, all outside the submitted work. All other authors declare no competing interests.

## Funding

This study has been supported by the Erasmus Trustfonds and the Erasmus Foundation.

## Authors’ contributions

K.J., W.S.G., H.J.G.W. and A.A.M.V. wrote the manuscript, which all authors have reviewed critically. W.S.G. and H.J.G.W. performed the bioinformatical analyses. K.J. and A.A.M.V. assessed the clinical data. G.J.L.H.L. reviewed and assessed the histopathological data. P.H., H.M.W., A.B., S.F.O, J.M.R., L.V.B., M.L. and R.H.J.M. are clinical contributors. M.P.L. and S.S. participated in the board of the CPCT-02 study. E.C. coordinated the sequencing of samples and contributed to the bioinformatical analyses.

## Acknowledgements

This publication and the underlying study have been made possible partly based on the data that Hartwig Medical Foundation and the Center of Personalized Cancer Treatment (CPCT) have made available for the study. This study has been made available by the Erasmus Trustfonds and the Erasmus Foundation. Furthermore, we would like to thank the nationwide network and registry of histo- and cytopathology in the Netherlands (PALGA) for their assistance in retrieving and storing the pathological records of the included patients with RCC. Figure 1 was created with BioRender (https://biorender.com/).

## Notes

### Summary of Updates

-

