## Supplementary material for "The genomic and transcriptomic landscape of advanced renal cell cancer for individualized treatment strategies": Legends supplementary figures

**Supplementary figure 1: Whole-genome sequencing and RNA-Seq metrics**

Panel **A** shows the genome-wide mean sequencing coverage for each sample (*N*=91) as boxplots, for both tumor (red) and whole-blood (blue). Panel **B** shows the number of mapped reads for the RNA-Seq samples (*N*=28).

**Supplementary figure 2: Mutational overview of the whole-genome sequenced advanced Renal Cell Carcionoma (RCC) cohort (*N* = 91)**
Track **A** shows the genome-wide and coding region frequency for three small mutational types (single nucleotide variant (SNV), multi nucleotide variant (MNV) and InDel). Track **B** shows the frequency of each type of transition and transversion including C to T in CpG context. Tracks **C**, **D** and **E** show boxplots of the tumor purity, the estimated genome-wide ploidy for the tumors and the ratio of transitions (Ti) to transversions (Tv). Track **F** shows the frequency in boxplots of six types of structural variants (translocation, deletion, tandem duplication, break-ends, insertion and inversion), and track **G** shows the consequences at coding level of the small mutational types. ccRCC = clear cell renal cell carcinoma. pRCC = papillary renal cell carcinoma. Undefined subtype = renal cell carcinoma, with undefined subtype. chRCC = chromophobe renal cell carcinoma. CDC = collecting duct carcinoma. tRCC = tubulocystic renal cell carcinoma.

**Supplementary figure 3: Relative contribution for COSMIC single base substitution mutational signatures across 100 bootstraps**
Panel **A** X-axis shows the relative contribution of the bootstrapped mutational signatures and the y-axis shows the SBS signature ordered by maximum relative contribution assigned across 100 bootstraps. Panel **B** X-axis shows all samples and the y-axis shows the relative contribution of SBS40 assigned across 100 bootstraps, with the red circle illustrating the originally assigned relative contribution, for comparison.

**Supplementary figure 4: Circos plots of chromothripsis samples from the Renal Cell Carcinoma WGS sequencing cohort**

Chromosomes indicated with a red block were flagged by chromothripsis detection. The star indicates the presence of a translocation from chromosome 3 to chromosome 5. Lines in the center indicate structural variants, with colors indicative of the type. ccRCC = clear cell renal cell carcinoma. pRCC = papillary renal cell carcinoma. Undefined subtype = renal cell carcinoma, with undefined subtype. chRCC = chromophobe renal cell carcinoma. CDC = collecting duct carcinoma. tRCC = tubulocystic renal cell carcinoma.

**Supplementary figure 5: Circos plots of non-chromothripsis samples from the Renal Cell Carcinoma WGS sequencing cohort**

Chromosomes indicated with a red block were flagged by chromothripsis detection. The stars indicates the presence of a translocation from chromosome 3 to chromosome 5. Lines in the center indicate structural variants, with colors indicative of the type. ccRCC = clear cell renal cell carcinoma. pRCC = papillary renal cell carcinoma. Undefined subtype = renal cell carcinoma, with undefined subtype. chRCC = chromophobe renal cell carcinoma. CDC = collecting duct carcinoma. tRCC = tubulocystic renal cell carcinoma.

**Supplementary figure 6: Clinical heterogeneity in Renal Cell Carcinoma WGS sequencing cohort**
Alluvial diagram from pretreatment status (systemic treatment before biopsy) to treatment (therapies received after biopsy) and on to RECIST score (v1.1 after first treatment). Colors indicative of RCC subtype.

**Supplementary Figure 7: Copy-number analysis in clear cell Renal Cell carcinoma (ccRCC)**
Panel **A** is a circular overview of the copy-number alterations in the WGS advanced ccRCC cohort (*N* = 72). The outer ring shows the chromosomal ideogram, followed by a cohort-wide GISTIC2.0 G-score track with large peaks rounded to 1 and -1. Negative copy-numbers on the y-axis (blue) indicate deletions, with positive (green) indicating amplifications. The darker color is indicative of passing the statistical *q*-value threshold of 0.05. Known cancer driver genes overlapping copy-number peaks found to be significant by GISTIC2.0 are labelled in the center of the circle, utilizing the same color scheme as the G-score track. Panel **B** displays the arm-level copy-number alterations of significantly altered (*q*-value < 0.05) chromosome arms according to GISTIC2.0 (>50% of arm affected) in the WGS cohort (*N* = 72). Each column represents an ccRCC sample (ordered descendingly by tumor mutational burden), with the chromosome arm listed on the y-axis. Increase in copy-number is displayed in the grid cells in yellow and decrease in purple, white squares show no change in copy-number at arm-level.

**Supplementary figure 8: *t-*distributed stochastic neighbor (*t*-SNE) embedding plot of the expression analysis from RNA sequencing samples from cohort of clear cell Renal Cell Carcinoma (ccRCC; *N* = 24) and papillary Renal Cell Carcinoma (pRCC; *N* = 4)**
CcRCC displayed in purple, pRCC in pink.

**Supplementary figure 9: Heatmap (A) and *t*-distributed stochastic neighbor embedding (*t*-SNE) plot (B) of samples from the Renal Cell Carcinoma RNA sequencing cohort**
Illustrative figures to visualize clustering of RCC samples with undefined subtype included. Gene selection and order in **A** is based on variance stabilized values with unsupervised clustering of Z-scores of the top 100 genes (based on smallest adjusted *p*-value) statistically significant genes, based on the differential expression analysis between ccRCC (*N* = 24) and pRCC (*N* = 4). **B** *t*-SNE plot of variance stabilized read counts of all protein coding genes in ccRCC, pRCC and RCC with undefined subtype.
