## Supplementary figures and images for "The genomic and transcriptomic landscape of advanced renal cell cancer for individualized treatment strategies"

### SuppFi9_UndefinedSubtype.pdf

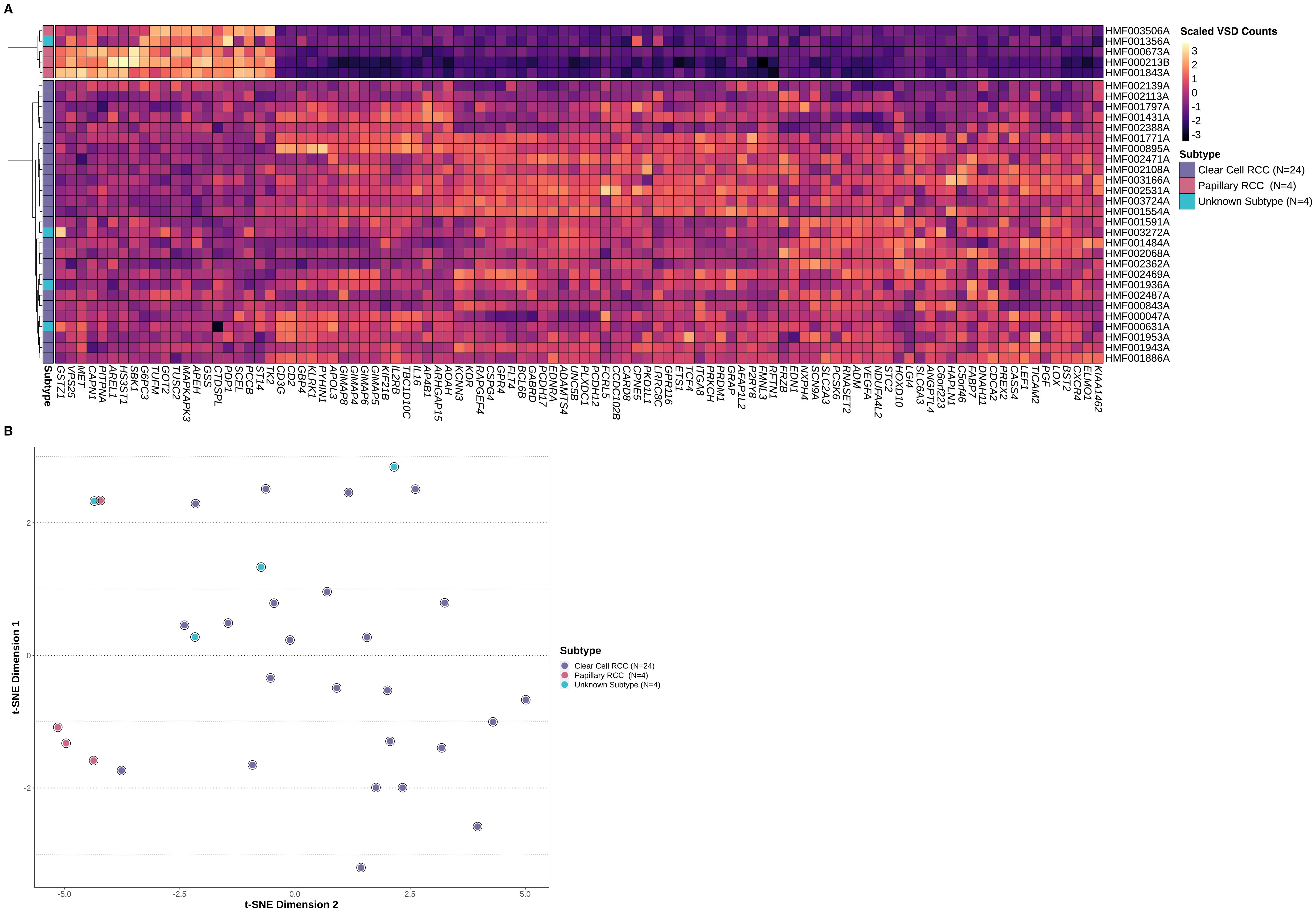

### SuppFig1_Metrics.pdf

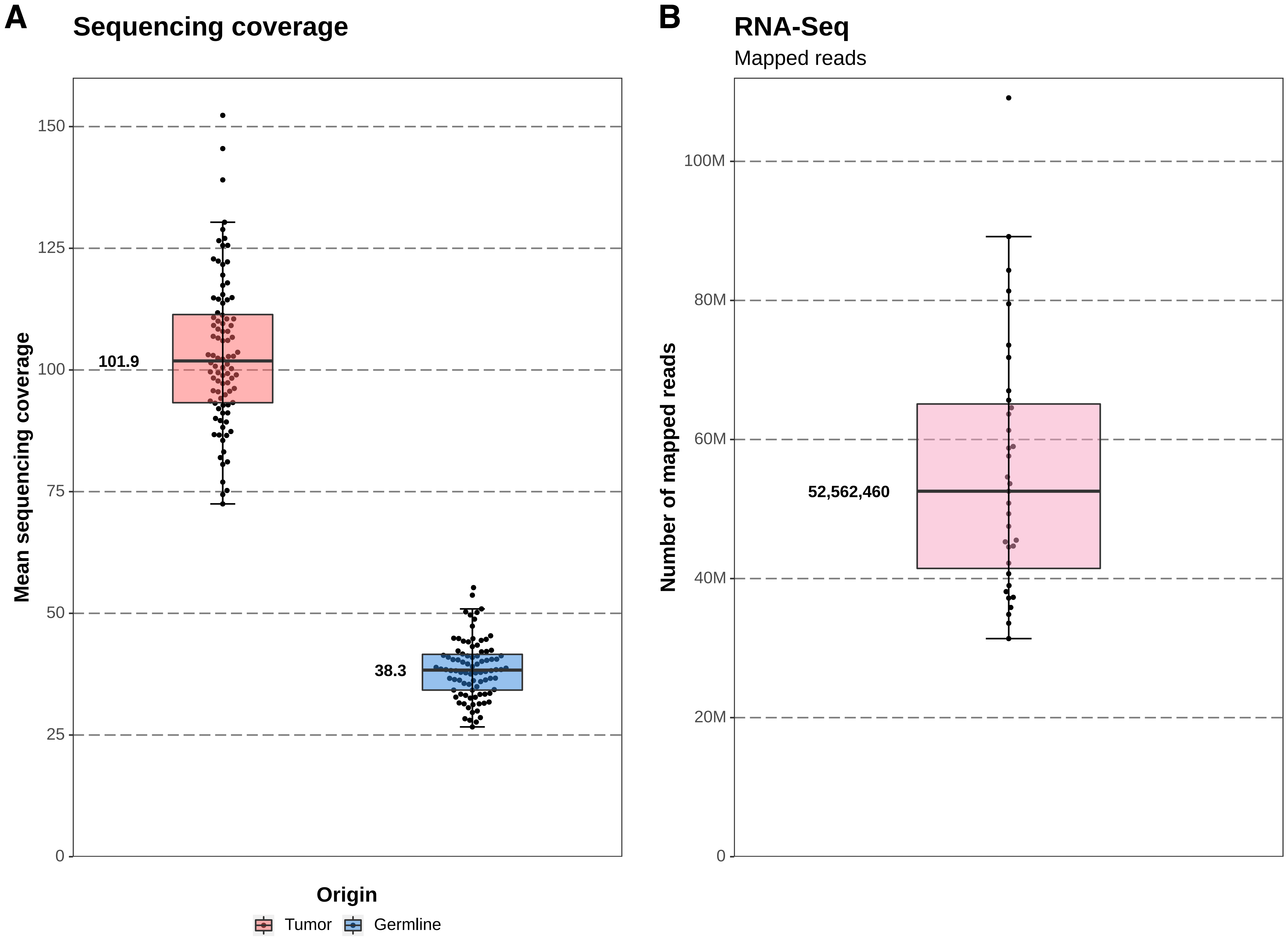

### SuppFig2_overview.pdf

#### Overview of Mutational Frequencies

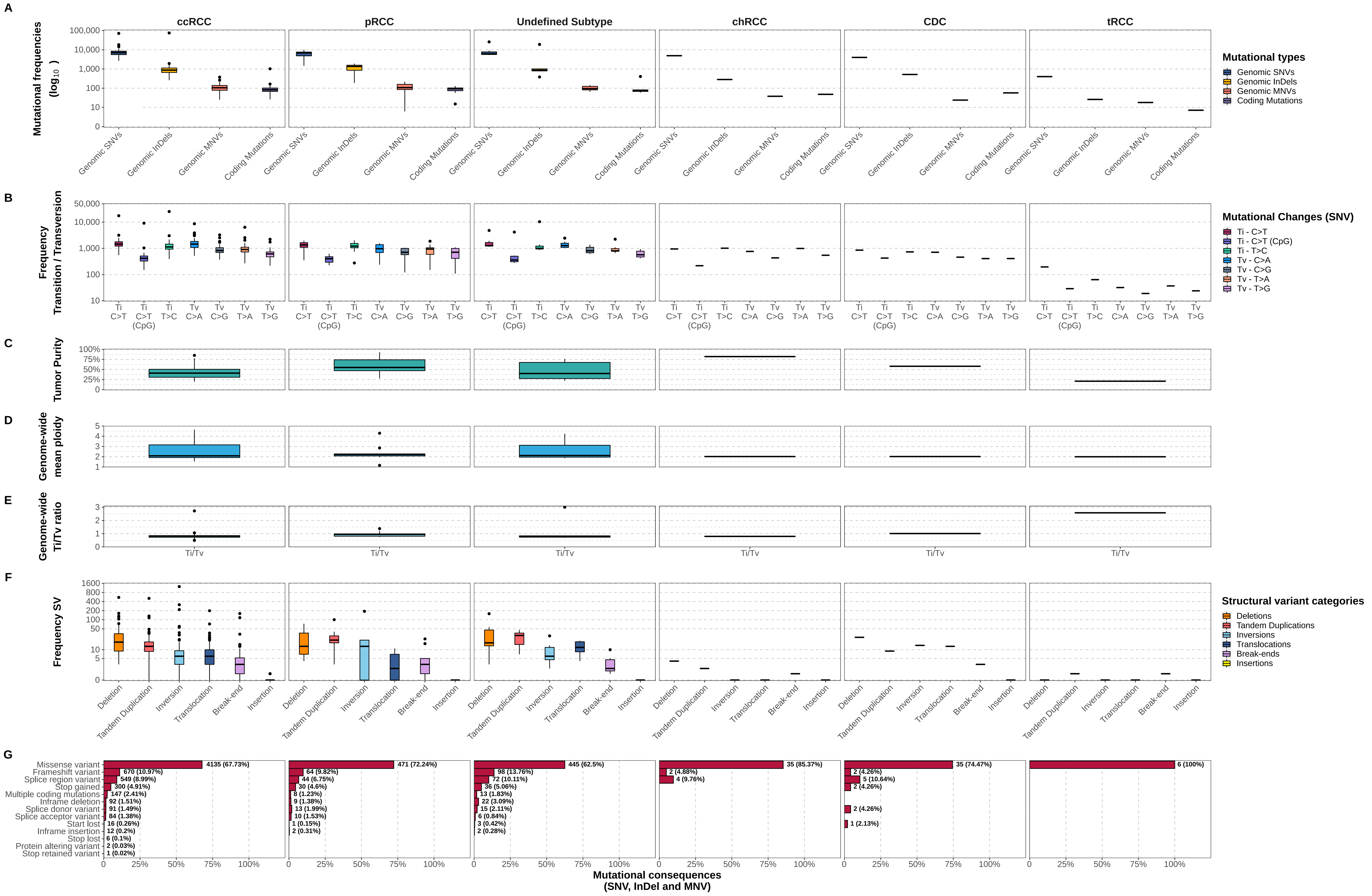

### SuppFig3_COSMIC.pdf

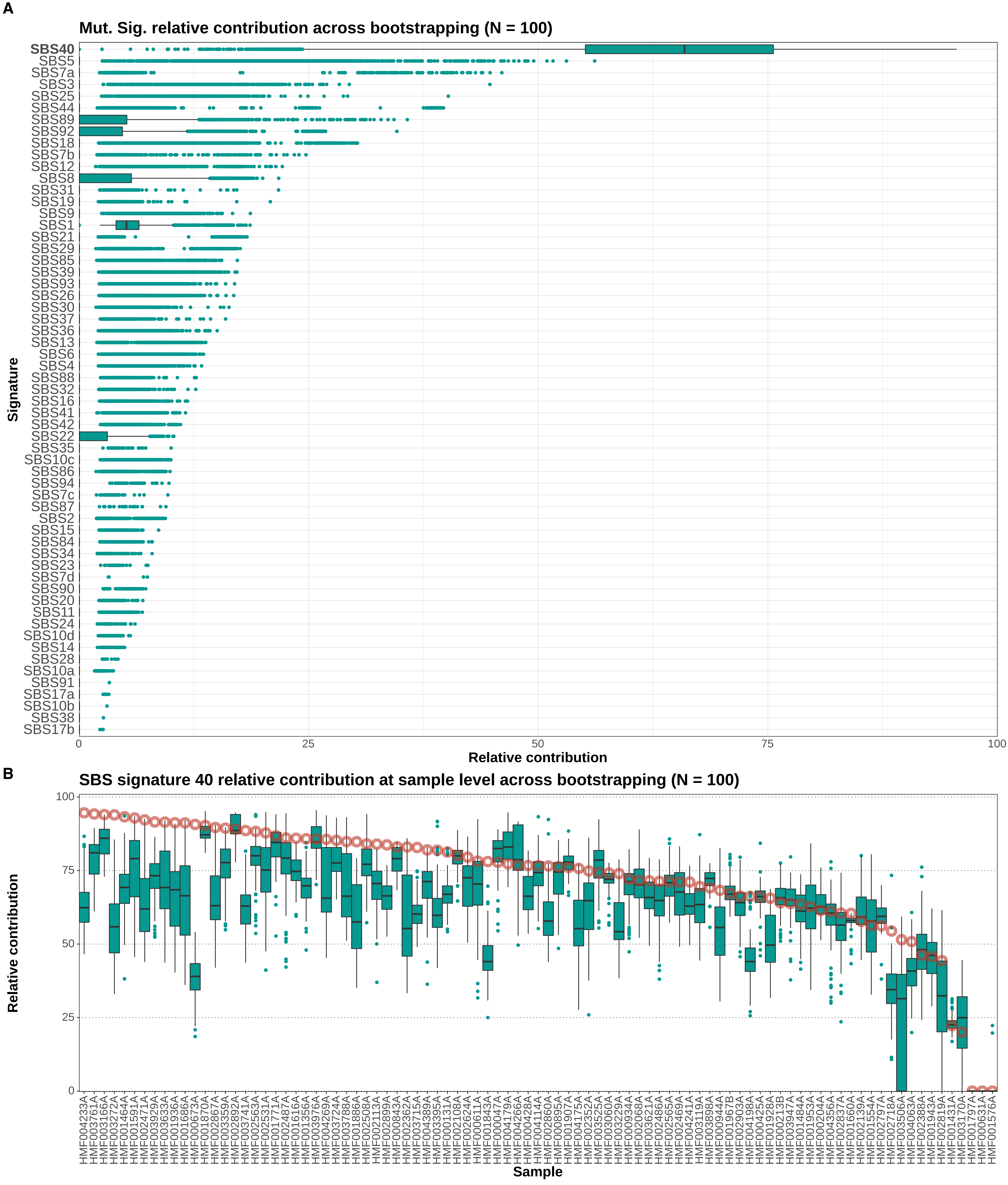

### SuppFig4_chromothripsis.pdf

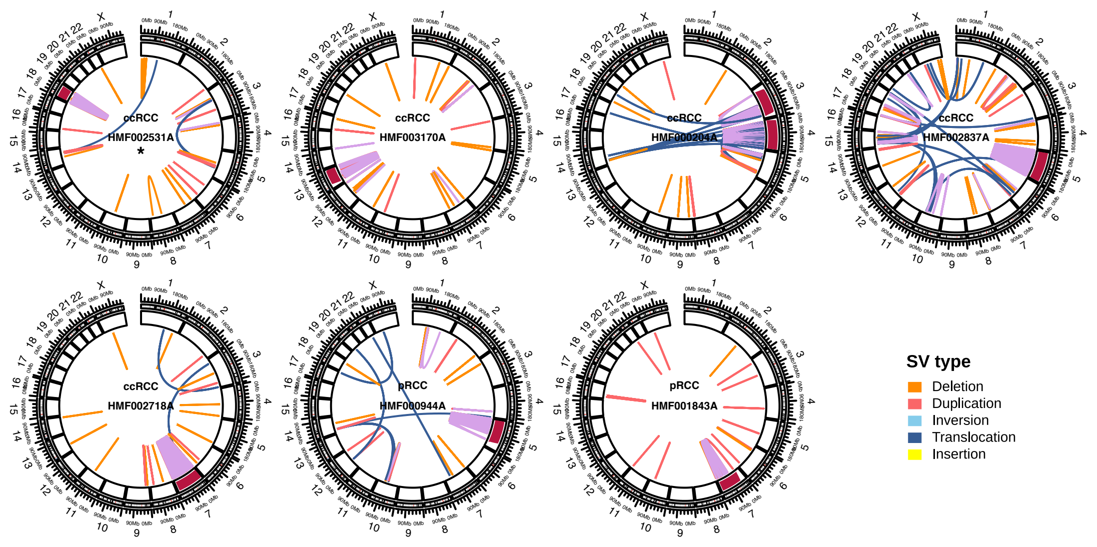

### SuppFig5_nonchromothripsis.pdf

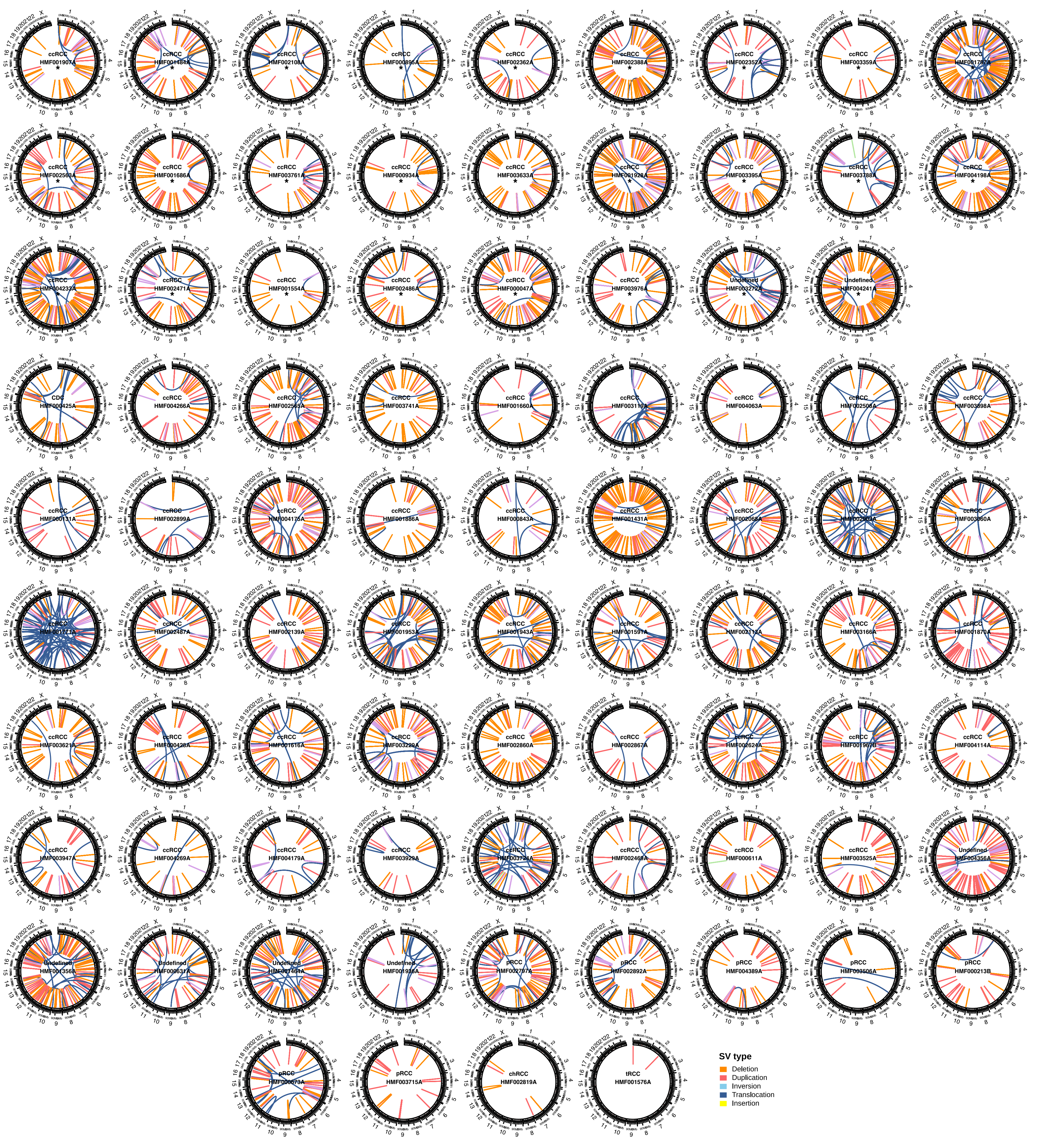

### SuppFig6_Heterogeneity.pdf

#### Supplementary Figure: Overview of heterogeneity

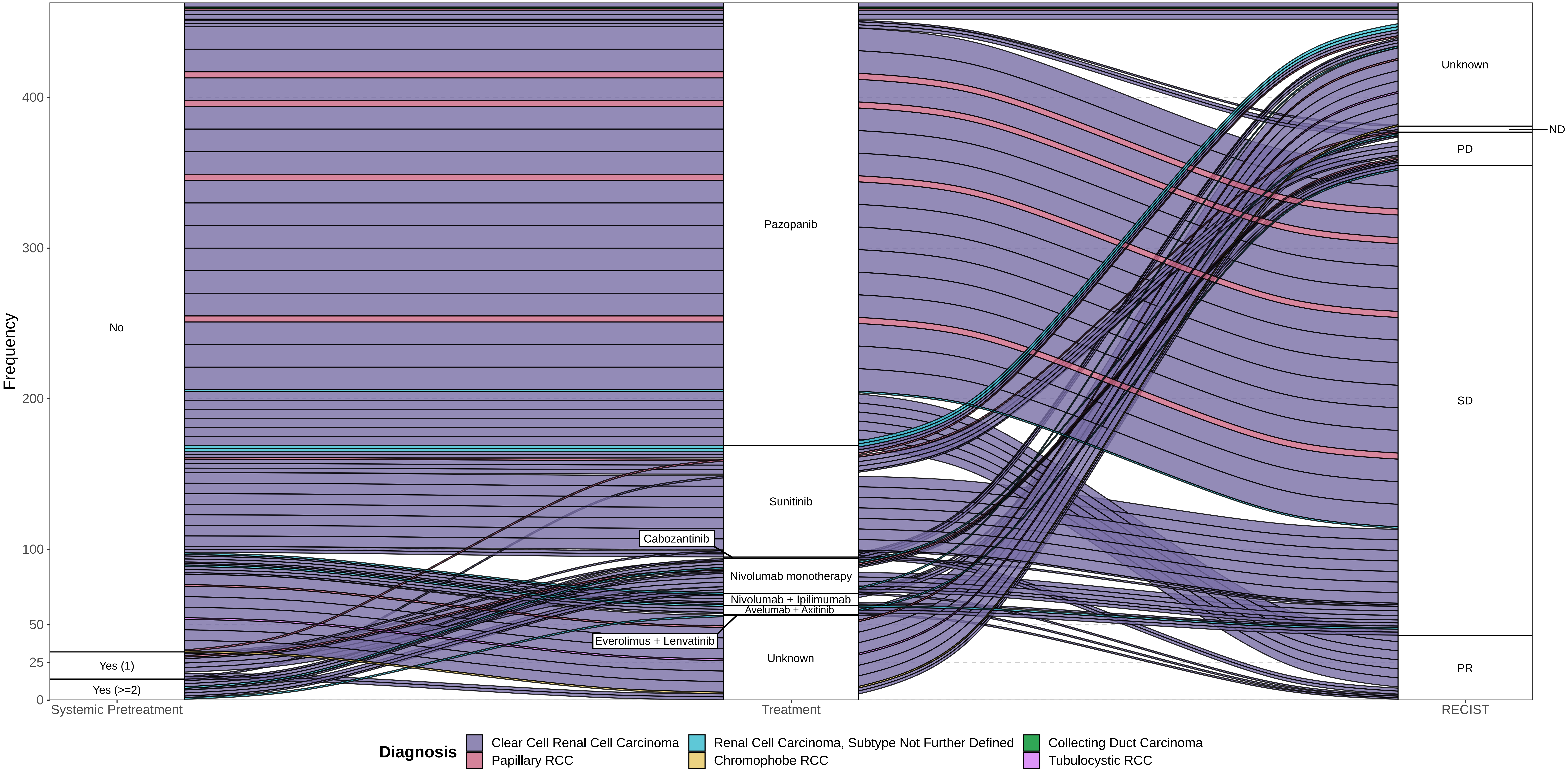

### SuppFig7_GISTIC.pdf

##### GISTIC2.0 analyses - Clear Cell Renal Cell Carcinoma

**A**

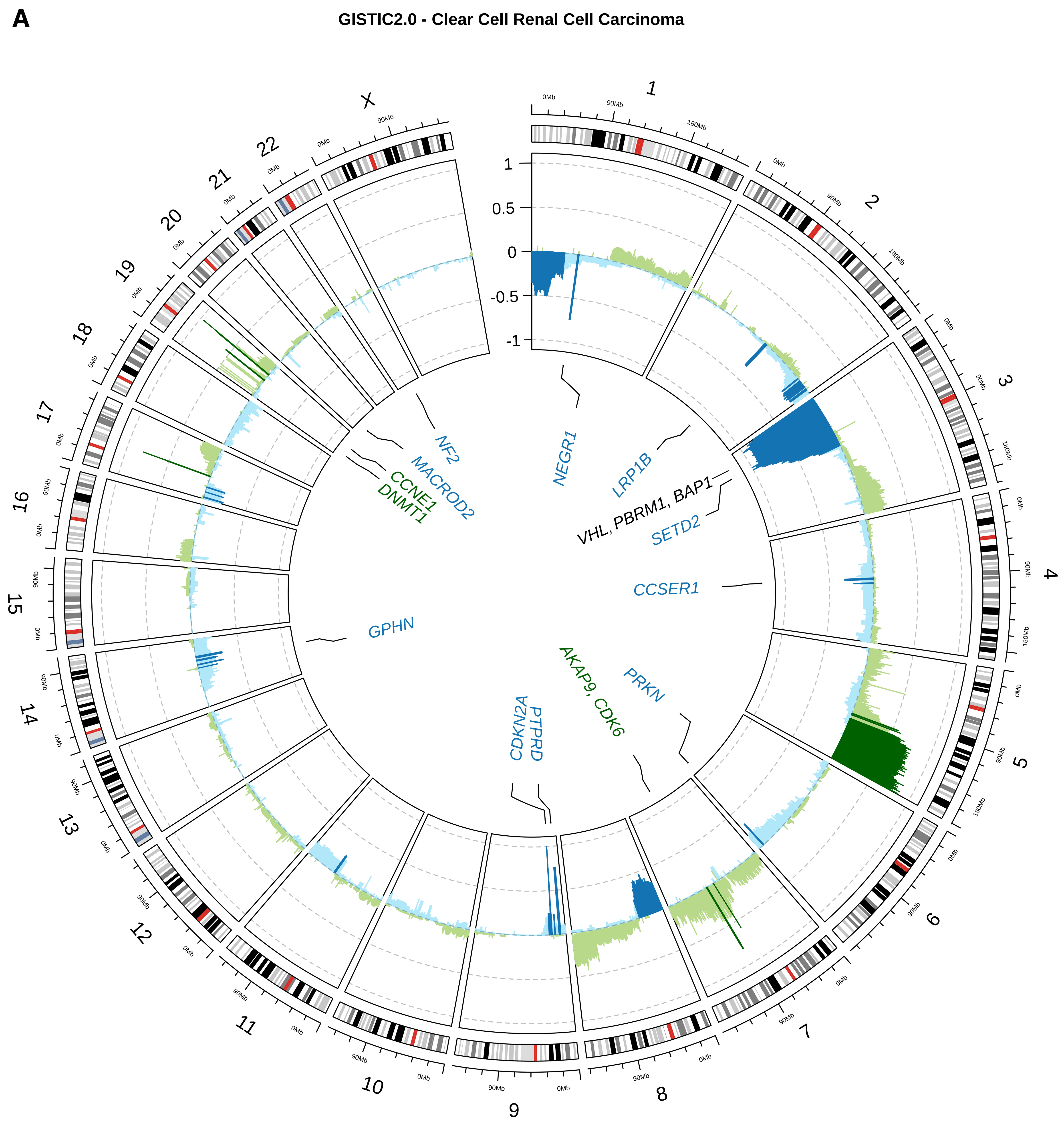


**B**

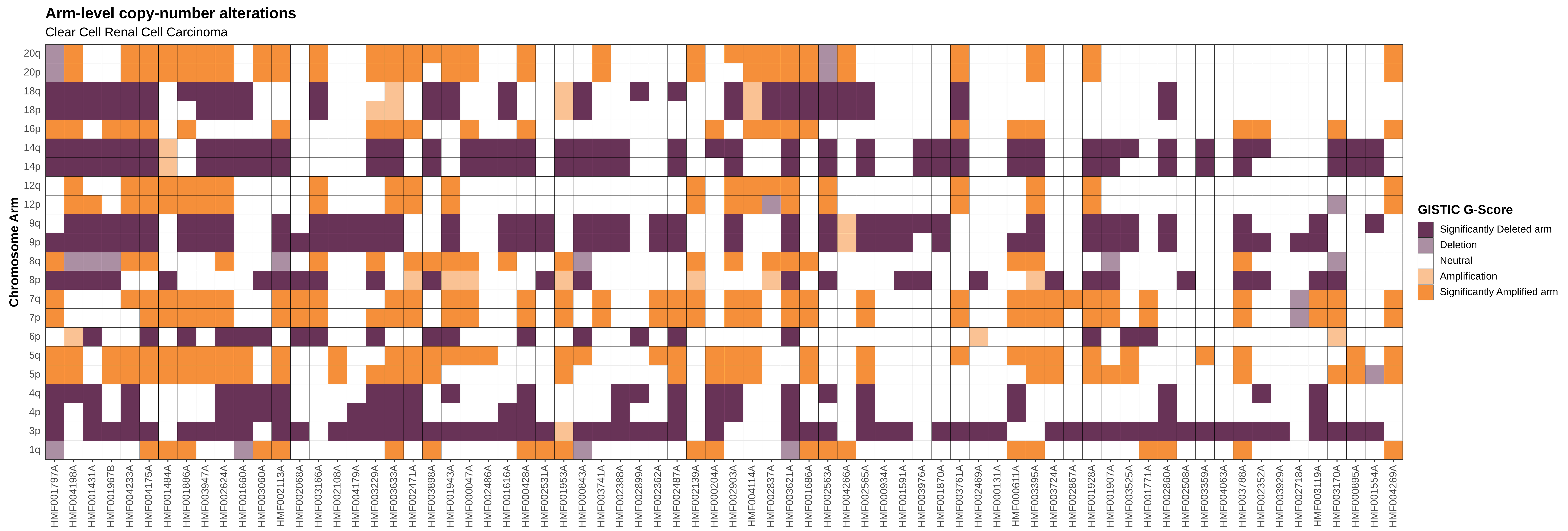

### SuppFig8_tSNE.pdf

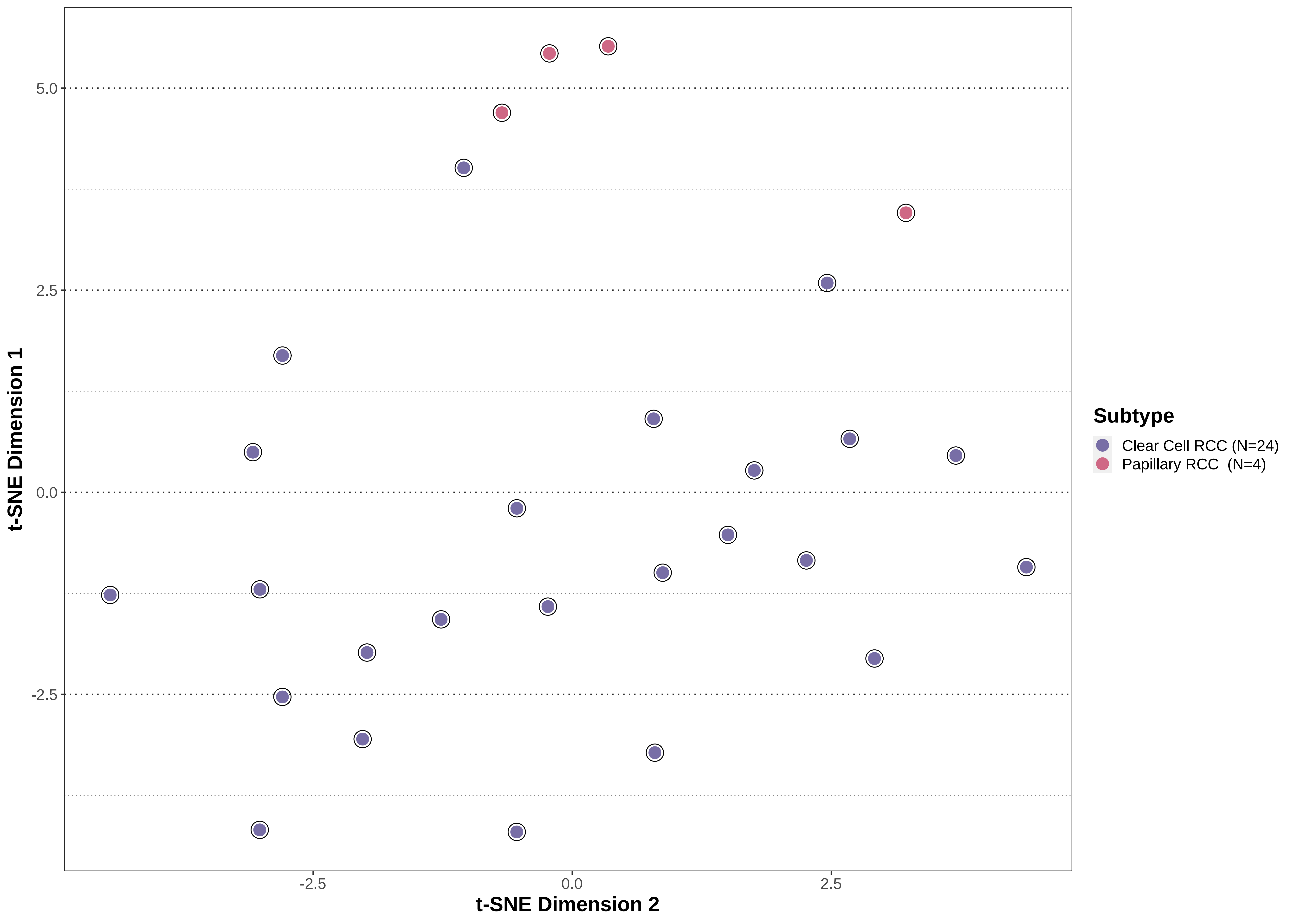
